# Prosocial behavior, social reward and affective state discrimination in adult male and female mice

**DOI:** 10.1101/2022.08.19.504492

**Authors:** Klaudia Misiołek, Marta Klimczak, Magdalena Chrószcz, Łukasz Szumiec, Anna Bryksa, Karolina Przyborowicz, Jan Rodriguez Parkitna, Zofia Harda

## Abstract

Prosocial behavior, defined as voluntary behavior intended to benefit another, has long been regarded as a primarily human characteristic. In recent years, it was reported that laboratory animals also favor prosocial choices in various experimental paradigms, thus demonstrating that prosocial behaviors are evolutionarily conserved. Here, we investigated prosocial choices in adult male and female C57BL/6 laboratory mice in a task where a subject mouse is equally rewarded for entering any of the two compartments of the experimental cage, but only entering of the compartment designated as “prosocial” rewards an interaction partner. In parallel we have also assessed two traits that are regarded as closely related to prosociality: sensitivity to social reward and the ability to recognize the affective state of another individual. We find that female, but not male, mice increased frequency of prosocial choices from pretest to test. At the same time, both sexes showed similar rewarding effects of social contact in the conditioned place preference test, and similarly, there was no effect of sex on affective state discrimination measured as the preference for interaction with a hungry or relieved mouse over a neutral animal. These observations bring interesting parallels to differences between sexes observed in humans, and are in line with reported higher propensity for prosocial behavior in human females, but differ with regard to sensitivity to social stimuli in males.

## Introduction

Prosocial behavior, defined as acting to meet the perceived need of another individual, is regarded as the highest form of empathy^1,2^. In humans, a major factor affecting the propensity for prosocial behaviors is gender^3^. It was observed that females have superior emotion discrimination abilities^4^, are more concerned about the well-being of others^5^, and utilize more resources to support others in need^6^. It was proposed that altruistic, prosocial behavior is a uniquely human characteristic (e.g.,^7^,^8^); however, a growing number of reports show that targeted helping is also observed in other species. Laboratory rodents (rats:^9–11^, mice:^12,13^, but see^14^) and some bird species^15^ prefer actions that reward another conspecific in choice tasks or free another animal from a restraint, with whom they share a food reward afterward. Unlike in humans, there are limited data on the effect of sex on prosocial behaviors in laboratory animals. Most rodent studies on affective state discrimination have focused on only one sex, usually males (for reviews, see rats&mice:^16,17^), although some studies have investigated females (e.g.,rats:^18–20^). Several studies examined both females and males, but the results considering sex-differences appear inconclusive (rats&mice:^21^, rats:^9,22,23^ mice:^24–26^). Some of the reports suggest that females are more susceptible to emotional contagion (mice:^25^), show enhanced emotion discrimination abilities in double approach paradigms (mice:^24^, rats:^27^) are more likely to perform prosocial actions (rats:^9,28^). However, other studies provide evidence for equal susceptibility of male and female rodents to emotional contagion (rat&mice:^21^, mice:^22,29–31^), equal affective state discrimination skills (mice:^26^), and comparable levels of prosocial behaviors (rats&mice:^21^, rats:^32^, prairie voles:^33^). Interestingly, some authors observed even higher levels of empathy-motivated behaviors in male rodents (rats&mice:^21^, rats:^23,34^, mice:^12,35^). Thus, previous reports appear inconsistent with regard to superior empathic and prosocial abilities in female mice.

Recently, it was proposed that prosocial behavior is directly motivated by the rewarding effects of social interactions as well as empathy, together termed the “camaraderie effect”^36^. This theory predicts that if sex differences in prosocial behavior were to be found between sexes of a given species, they should also be found in social reward and empathy, or its prerequisite, affective state discrimination. If proven correct, the theory would potentially provide a framework to at least partly reconcile reported differences in prosocial behaviors in mice^12–14,35^. Thus, we first investigated prosocial choices in adult male and female C57BL/6 laboratory mice, towards a familiar partner (sibling) using model based on the general outline the rat task described by Hernandez-Lallement and collaborators^10^. Affective state discrimination was tested in a mice paradigm modified from Scheggia et al. in 2020, where sensitivity to the emotional state of interaction partners was assayed^26^ and social reward was tested in the social conditioned place preference task^37^. We found that, similar to humans, sex had a significant effect on the propensity for prosocial choices; however, females and males did not differ with respect to levels of social reward and affective state discrimination.

## Results

### Prosocial choices in adult male and female mice

To assess prosocial choices in adult mice, we used a custom-made maze, as shown in **Figure 1a**. In the task, a focal animal, the actor, chose to enter one of two compartments and would be rewarded with chocolate chips for either choice. A second animal, the partner, also received a reward, but only if the actor entered the compartment designated “prosocial”. The wall separating the actor’s and partner’s compartments was transparent and perforated, allowing for visual, auditory and olfactory communication. The schedule of the experiment is summarized in **Figure 1b**. First, the actor animals underwent up to 4 pretest sessions, one session per day, without partner animal (**Fig. 1c**). The number of chocolate chips consumed was checked after each trial, and only animals that consumed at least 85% of the pellets over two consecutive days were subjected to the actual test (**Fig. 1d**). The average number of sessions required to reach this criterion was 2.4 and 2.63 in male (n=10) and female (n=8) mice, respectively (t-test, t_16_ = 0.76, p = 0.46, **Fig. S1a-d**, **Table S1**). The pretest was intended to train actors and to assess their inherent preference between the left and right compartments in the absence of a partner animal. No inherent preference for cage side was detected in males or females (Fig. S1e, Table S2).

**Figure 1.**
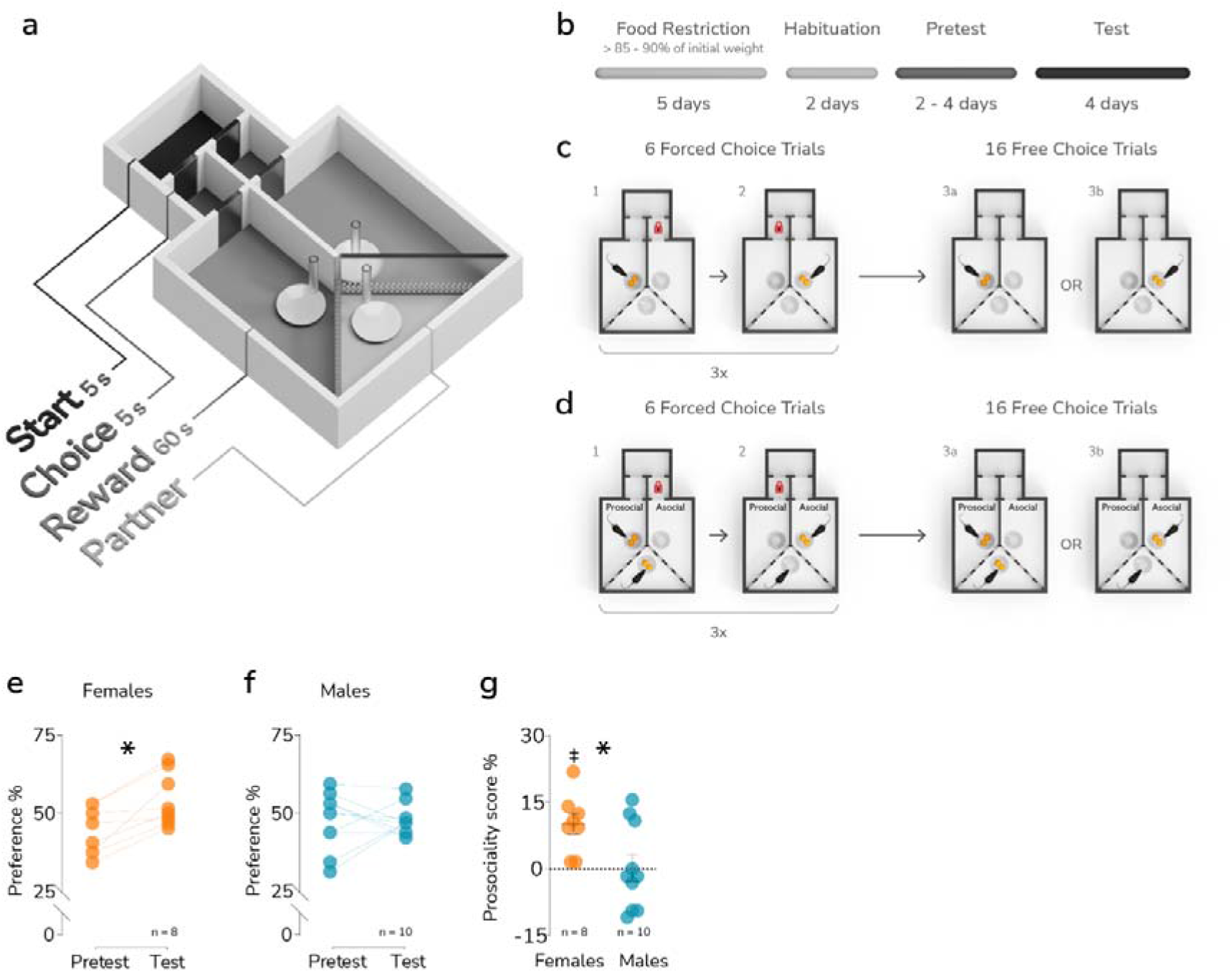
Prosocial choices in male and female mice. (**a**) A schematic representation of the testing apparatus. (**b**) Experimental schedule. (**C-D**) A diagram summarizing the pretest and test phases of the experiment, respectively. (**e-f**) The change in preference for the ‘prosocial’ compartment between the pretest and test phases in female and male mice, respectively. The results shown are the mean values from all sessions in the pretest and test phases. A significant difference between the phases is marked with a “*” (paired t-test p < 0.05). Respective group sizes are indicated below the graphs. (**g**) The difference between females and males prosociality score, calculated as test preference for the prosocial compartment minus pretest preference for the same compartment (%). The bar and whiskers represent the mean and s.e.m. values. A significant difference between means is marked with a “*” (t-test p<0.05), and a significant difference vs. 0 is indicated by a “‡” (one-sample t test p<0.05). The group sizes are indicated below the graph.

Then, the partners were introduced, and one of the compartments was designated “prosocial” (**Fig. 1d**). Four test sessions were performed. The frequency of entries to the prosocial compartment during the test was compared to the frequency of entries to the same compartment (“prosocial to be”) during the pretest. In female mice, the preference for the prosocial compartment increased significantly (**Fig. 1e**, **Table S2**), while male animals appeared to show no change from their initial choices (**Fig. 1f**, **Table S2**). In the case of females, the preference for prosocial behavior changed from 44.14% initially to 54.30% (average from the 4 trials, paired t-test, t_7_ = 4.33, p = 0.003), while in males, these values were 47.52% and 47.83%, respectively. The difference in the prosociality score (defined as the percentage of prosocial choices during the test minus the percentage of prosocial choices during the pretest) between males and females was significant (**Fig. 1g**, **Table S3**, t test, t_16_ = 2.47, p = 0.025). Additionally, we examined correlations between the absolute and relative weights of the actor and partner and the prosociality score. Relative weight was defined as the difference in weight between actor and partner calculated as a percentage of actor’s weight. The analysis revealed no significant association for absolute weights in any of the sexes and no significant correlation of relative weight in the case of females (**Table S4**). However, a negative correlation between relative weight and prosociality score was observed in male mice (**Table S4**, r = −0.73, p = 0.014). Thus, we found that female mice favored prosocial choices in the task, while males have shown no preference for either choice.

### Social reward

An increase in the frequency of prosocial choices observed in females is evidence of reinforcing effects of their consequences and, thus, of a rewarding effect of the choice. No preference for the prosocial choices in male mice could potentially be attributed to a generally lower sensitivity to the rewarding effects of social interaction. To assess this possibility, we tested adult mice of both sexes in the social conditioned place preference task (sCPP), with a 6-day conditioning protocol^37^. In this test, experience of social contact during conditioning causes an increase in time spent in the previously neutral context from pretest to posttest. Both female (n=16) and male (n=12) mice significantly increased the time spent in the context associated with group housing (**Fig. 2a**, paired t-test, t_15_ = 2.825, p = 0.012, **Fig. 2b**, t-test, t_11_ = 4.202, p = 0.002, **Table S5**). Likewise, the preference score (i.e. the difference in time spent in the social minus isolate context during the posttest) was significantly higher than chance value in both female (one sample t-test, t_15_ = 3.282, p = 0.005) and male (t-test, t_11_ = 2.446, p = 0.033) mice (**Fig. 2c**, **Table S6**). Moreover, there was no difference between males and females in preference score (t-test, t_26_ = 0.5, p = 0.62). These results indicate that social interactions with siblings were rewarding for male and female mice to similar extent. The finding that sCPP can be elicited in adult mice using contact with age and sex matched conspecifics as a reward appears contradictory to the previous findings by Nardou and collaborators in 2019, who have shown that sCPP was no longer observed at the age of 8 weeks in male, and at 11 weeks in female mice^38^. However, in the study the short (2 days) conditioning protocol was used^38^, and we have recently shown that sCPP can be observed in adult (>11 weeks) female mice, when the longer, 6-day, conditioning protocol is used^37^. Here, to assess if the effect of conditioning length is similar in males, we performed additional experiment on male mice (n=8) using a 2-day conditioning protocol (**Fig.2d-e**, **Table S5**–**S6**). We found no effect of social conditioning, which is in agreement with the results reported in 2019 by Nardou and collaborators^38^.

**Figure 2.**
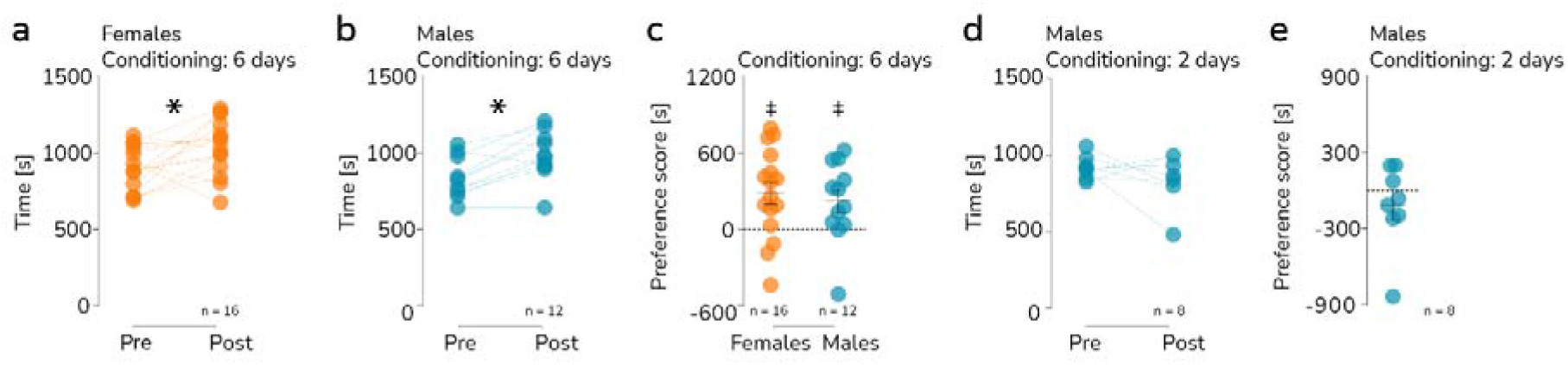
Social conditioned place preference. (**a-b**) The change in preference for the social context after 6 days of conditioning in female and male mice, respectively. Each pair of points represents an individual animal, and the group sizes are indicated below the graphs. A significant difference between the pre- and posttests (30 min) is marked with a “*” (paired t-test, p<0.05). (**c**) No difference between females and males preference score [s], calculated as time spent in social context posttest [s] minus time spent in in isolate context posttest [s]. Each point represents an individual female or male mouse, with the mean and s.e.m. shown in black and the group sizes indicated below. A significant preference (greater than 0) is indicated by a “‡” (one-sample t-test p<0.05). (**d**) No significant change in preference for the social context after 2 days of conditioning in male mice. (**e**) The preference score for social context during the posttest after 2 days of conditioning. The group sizes are indicated below the graph.

### Sensitivity to the affective state of the interaction partner

A plausible explanation for the observed effect of sex on prosocial choices would be a difference in sensitivity to social cues during interaction. To test this possibility, mice were assayed for their preference for interaction with a “neutral” vs. altered affective state demonstrator (“relieved” or “hungry”) in a procedure based on the task described in 2020 by Scheggia et al.^26^ and summarized in **Figure 3a**. During the main phase of the test, the animal tested (observer) was placed in the cage where demonstrators were present, both confined under wire cups. One of the demonstrators was food deprived: for 24h (hungry) or for 23 hours and then offered *ad libitum* food access (standard laboratory chow) for 1 h preceding the test (relieved). The second demonstrator (neutral) as well as the observer had constant *ad libitum* food access. Both female (n=12) and male (n=13) observer mice spent significantly more time sniffing the relieved demonstrator during the first minute of the test (**Fig. 3b-c**, **Table S7**; **Fig. 3b**, repeated measures ANOVA, effect of time, F_(2.94, 64.62)_=3.69, p = 0.017, Šídák’s test, sniffing relieved vs. neutral demonstrator during first minute, p = 0.019; **Fig. 3c**, effect of demonstrator’s state, F_(1, 24)_ = 6.233, p = 0.012, Šídák’s test, sniffing relieved vs. neutral demonstrator during first minute, p = 0.02), and there was no effect of sex on the fraction of time spent sniffing the relieved demonstrator (**Fig. 3d**, **Table S8-9**). The preference for relieved demonstrator was most evident during the first minute of the observation, which corroborates the results obtained earlier^26^. Thus, only the behavior during this time period was considered an indication of affective state discrimination. Similarly, both female (n=8) and male (n=10) observer mice spent significantly more time sniffing the hungry demonstrator during the first minute of the test (**Fig. 3e-f**, **Table S7**; **Fig. 3e**, repeated measures ANOVA, effect of interaction of time x demonstrator’s state, F_(3, 42)_ = 5.157, p = 0.004, Šídák’s test, sniffing hungry vs. neutral demonstrator during first minute, p = 0.008; **Fig. 3f**, effect of interaction of time x demonstrator’s state, F_(3, 54)_ = 5.391, p = 0,003, Šídák’s test, sniffing hungry vs. neutral demonstrator during first minute, p = 0.026), and there was also no effect of sex (**Fig. 3g**, **Table S8-9**). The position of the altered affective state demonstrator (east vs. west side of the testing apparatus) was selected randomly, and no effect of the relieved or hungry demonstrator’s position on the percentage of time spent sniffing this demonstrator was found (**Fig. S2a-b**, **Fig. S2f-g**, **Table S10**). Additionally, to control for inherent preference of the position of the cups (east vs. west), we also analyzed the time spent sniffing the cup in which the relieved/hungry demonstrator would be placed during the last day of adaptation. The time spent sniffing the empty cup did not differ from chance level (**Fig. S2e**, **Fig. S2h**, **Table S11**) except for females paired later with relieved demonstrators. In this case time spent sniffing the empty cup was significantly shorter than the chance level (**Fig. S2e**, t-test, t_8_ = 2.495, p = 0.037). Finally, we also analyzed the correlations between the fraction of time spent sniffing the relieved/hungry demonstrator and the weight of the animals, both observers and demonstrators (as well as the weight difference between them). This analysis revealed a significant negative correlation for both relieved demonstrator (r = −0.7, p = 0.001) and observer weight (r = −0.62, p = 0.028), as well as a trend towards observer-relieved demonstrator weight difference (r = 0.53, p = 0.073), in females (**Table S4**). These results could indicate greater ability to recognize “relief” state among smaller females. No significant correlations were found in the case of hungry demonstrators, in either sex (**Table S4**). Taken together, these results show that both female and male mice discriminate between affective states of familiar conspecifics. Thus, there is no evidence of an effect of sex on sensitivity to affective states of conspecifics.

**Figure 3.**
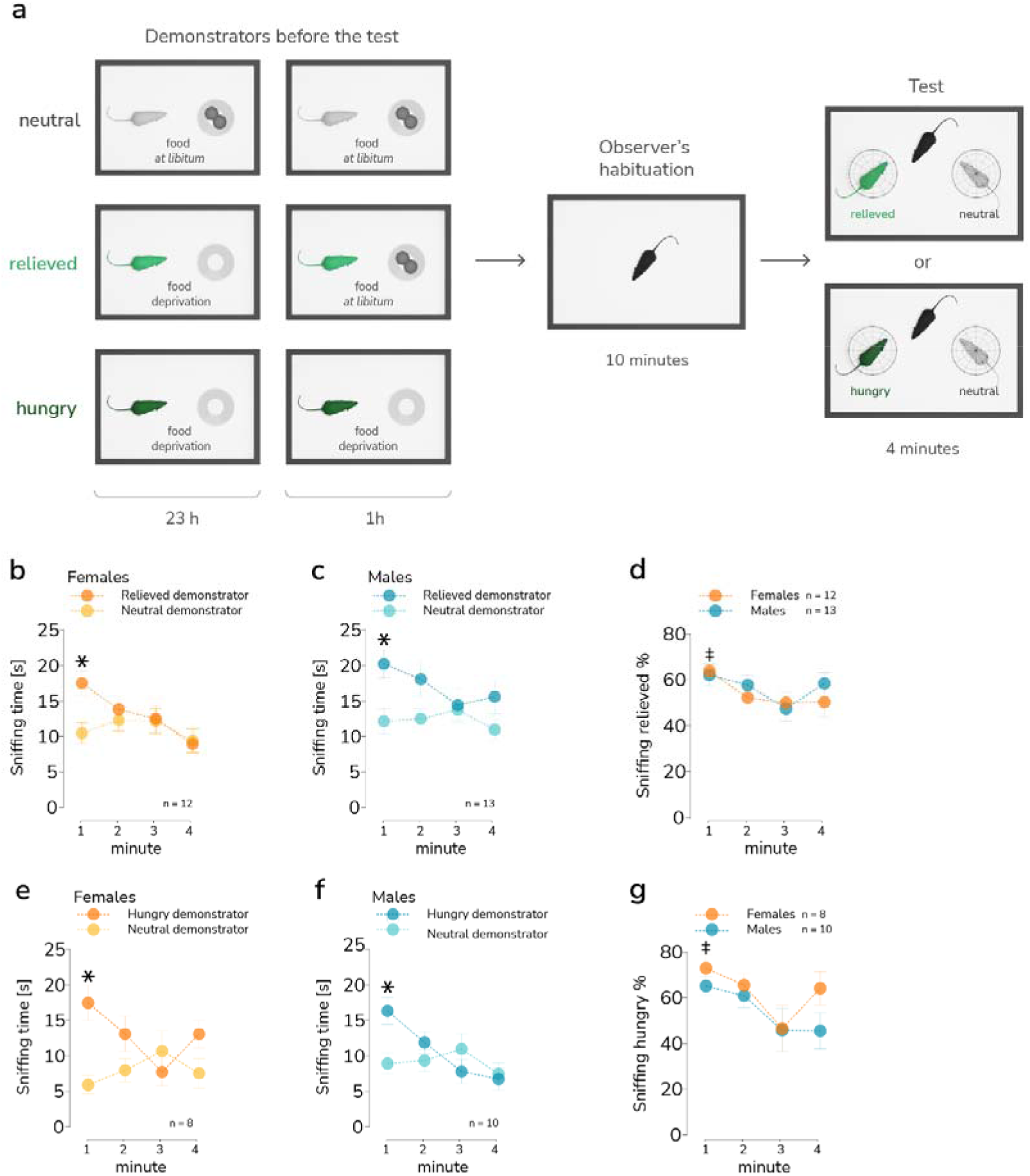
Affective state discrimination. (**a**) A schematic representation of the task. (**b-c**) The time spent by the demonstrator sniffing the relieved (darker points) and neutral (lighter points) demonstrators, female and male mice, respectively. Each point represents the mean time spent sniffing respective partners during a 1-minute interval (bin). The whiskers represent s.e.m. values, and significant differences between the mean time spent sniffing in the first one-minute interval are shown with a “*” (ANOVA with repeated measures; post hoc Šídák’s multiple comparisons test, p < 0.05). (**d**) No difference between female and male mice in preference for sniffing the relieved demonstrator over the neutral demonstrator. The points represent mean values over 1-minute intervals, whiskers represent the s.e.m. Significant differences vs. 50% during the first 1-minute interval in both female and male mice are indicated by a “‡” (one-sample t-test p<0.05). (**e-f**) The time spent by the demonstrator sniffing the hungry (darker points) and neutral (lighter points) female and male demonstrators, respectively. Each point represents the mean time spent sniffing respective partners during a 1-minute interval (bin). The whiskers represent s.e.m. values, and significant differences between the mean time spent sniffing in the first one-minute interval are shown with a “*” (ANOVA with repeated measures; post hoc Šídák’s multiple comparisons test, p < 0.05). (**g**) No difference between female and male mice in preference for sniffing the hungry demonstrator over the neutral demonstrator. The points represent mean values over 1-minute intervals, whiskers represent the s.e.m. Significant differences vs. 50% during the first 1-minute interval in both female and male mice are indicated by a “‡” (one-sample t-test p<0.05). The group sizes are indicated below the graph.

## Discussion

We found that female, but not male, C57BL/6 mice showed significant preference for prosocial behavior toward a familiar partner. Recently, Scheggia and colleagues showed that altruistic, prosocial behaviors in mice are dependent on sex, familarity, social hierarchy, and also internal and affective state^12^. They observed that male mice were more likely to share food with same sex conspecific than females in the operant two-choice social decision-making task, which is at odds with our results. This discrepancy may emerge from differences in methodology. In the report cited, testing was performed in an automated operant chamber, whereas we have used a manually operated cage. This could have affected animal’s stress levels and thus bias choices. Furthermore, Scheggia and collaborators found that tactile social contact is necessary for prosocial choice preference development in male mice^12^. When partition dividing actor and partner lacked perforations, the focal animals made fewer prosocial decisions. In our experiment transparent and perforated partition between actor’s and partner’s compartments was used, to allow access of visual, auditory and olfactory cues. However, due to the small size of the perforations, direct contact between mice was restricted, which speculatively could influence preference of prosocial choice in males. Our findings, together with the results reported by Scheggia and colleagues^12^, suggest that lower level of tactile contact might decrease prosocial behavior in male, but increase in female mice. Indeed, there is evidence for an effect of sex on processing of tactile contact. Experiments performed on rats showed that regular-spiking neurons in the barrel cortex exhibited stronger responses to facial touch (nose-to-nose) in males compared to females^39^. Future studies should directly test the relationships between sex, social touch and prosocial behavior.

It should be noted that the observation that female mice are more prosocial than males in a food-motivated prosocial choice task is in accordance with other findings in rats using different measures of prosociality. In 2011 Ben-Ami Bartal and colleagues found that females are more likely to learn how to free a trapped cagemate, and, when the task is learned, females perform it faster than males^9^. Furthermore, in 2020 Heinla and colleagues found that female, but not male, rats show consolation-like behavior directed toward cagemates that were recently attacked by another individual^28^.

Based on the “camaraderie effect” theory^36^, we hypothesized that a female advantage in prosocial behavior may stem from higher emotion discrimination abilities or higher rewarding effects of female–female, compared with male–male, social interactions. However, no sex difference in these behaviors was observed. The finding that emotion discrimination abilities, independently from valence, are similar in male and female mice corroborates previous observations Scheggia and colleagues^26^, nevertheless, some of the previous reports showed higher emotion discrimination abilities in female rodents (mice:^24^, rats:^27^). These discrepancies may stem from the differences in the severity of the emotion-eliciting stimulus and/or emotion valence. In our study, a positive state was induced by deprivation and subsequent provision of food and the negative state was induced by food deprivation for 24 h before the test, which could be considered as a relatively mild stress. Scheggia and collaborators used a test where a positive state was induced by deprivation and subsequent provision of water and the negative state was induced by 15 minutes of restraint before the test, which also is arguably a mild stressor. Conversely, studies that have demonstrated higher emotion discrimination abilities in female rodents used severe stimuli, i.e. pain or footshock (mice:^24^, rats:^27^). A conjecture that sex differences in affective state discrimination are evident only in the test involving highly stressful stimuli would be in agreement with “fitness threat hypothesis”, which states that female advantage in emotion recognition may be limited to negative emotional expressions, as they signal a potential threat to the offspring^40^.

The finding that rewarding effects of social interactions are similar in adult male and female mice is especially surprising, as males of the *Mus musculus* species studied in natural or seminatural conditions usually have been found to form territories and aggressively defend them from other males^41^. Female mice, in contrast, are capable of communal nesting and nursing^42^. Both male and female mice disperse from their natal groups, but males do this more frequently and at younger ages^43^. Taken together, these literature data suggest that the motivation for the social context preference observed here may differ between males and females. In females, amicable social interactions are the most likely cause of the increase in the preference for social context. In males, however, the opportunity for aggressive encounters may have caused the same effects. Earlier studies support this interpretation, as rewarding effects of aggression were repeatedly documented in male mice and rats (for a review, see^44^), but were absent in female mice^45^.

A potential limitation of our study is that in the affective state discrimination test the food deprivation was used instead of water deprivation, as previously described^26^. This leads to a question if observer mice indeed have shown preference for the emotionally aroused conspecific or rather the one that emitted more intense food odor. However, this issue is mostly resolved by the comparison of “relieved” and “hungry” conditions. If, in the “relieved” condition, relieved demonstrators emitted more smell of the food than neutral demonstrators, then, consequently, in the “hungry” condition, neutral demonstrators would have emitted more intense food odor than hungry demonstrators. If observer mice were attracted to the smell of food rather than the emotional arousal of the demonstrators, in the “hungry” condition they would have explored the neutral demonstrator more than a hungry one, which was not the case. Hence, we believe that in our version of the task mice indeed show recognition of emotional states of others, not interest in food odor. Taken together, our results show that, similar to humans, female mice tend to be more prosocial than males, but this difference may not stem from superior empathic abilities or higher rewarding effects of social interactions. Thus, the relationships among prosociality, affective state discrimination and social reward should be reconsidered, and correlations between these traits are not indicative of causation.

## Materials and methods

### Animals

Experiments were performed on C57BL/6 mice bred at the Maj Institute of Pharmacology Animal Facility. Mice were housed in a 12-hr light-dark cycle (lights on at 7 AM CET/CEST) under the controlled conditions of 22 ± 2°C temperature and 40-60% humidity. In the prosocial choice test, mice were housed as sibling pairs. For affective state discrimination, sCPP mice were housed with littermates of all the same sex or alone, depending on the phase of the experiment. Water was available *ad libitum*. Home cages contained nesting material and aspen gnawing blocks. Behavioral tests were conducted during the light phase under dim illumination (5-10 lux). Affective state discrimination and sCPP tests were video recorded with additional infrared LED illumination. The age and weight of all experimental groups are summarized in **Table S12**.

All behavioral procedures were approved by the II Local Bioethics Committee in Krakow (permit numbers 224/2016, 34/2019, 35/2019, 32/2021) and performed in accordance with the Directive 2010/63/EU of the European Parliament and of the Council of 22 September 2010 on the protection of animals used for scientific purposes. The reporting in the manuscript follows the ARRIVE guidelines.

### Prosocial choice test

The method was partly based on the prosocial test for rats described in 2014 by Hernandez-Lallement and colleagues^10^. The custom cage (main compartment: 30 × 36 × 30 cm; start-choice compartment: 12 × 14 × 30 cm) used in the procedure is shown in **Figure 1a**. The walls between reward and partner compartments were transparent and perforated so animals could see, hear, and smell each other during the experiment. Two pairs of doors were used for each arm of the apparatus to prevent the animal from going back to the starting compartment. Animals were given limited time to enter the choice compartment (5s) and reward compartments (5s). In case mice didn’t make a choice in time, animals were gently touched by hand of the experimenter to encourage them to enter either of the reward compartments. Entering one of two reward compartments resulted in food delivery after 5 seconds. The time to consume the reward was also limited (60s). Animals were moved to starting compartment if they consumed both rewards or 60s had passed. Time between each trial lasted 10s. The primary difference from the previously described apparatus is the single compartment for the interaction partner. The experimental schedule is summarized in **Figure 1b** and consisted of 4 phases: food restriction (5-7 days), habituation (2 days), pretest (2-4 days, depending on completion criterion; see **Table S1**), and main test (4 days). Mice had restricted access to food throughout the experiment. Habituation was performed when animals reached 85-90% of their initial body weight. On the last day of food restriction, the heavier mouse from each cage was selected as the actor, and the lighter mouse was used as the partner. The rationale was to increase the chance of observing prosocial behavior in actors, as it was shown that the number of reward portions provided to the hungry partner is negatively correlated with the partner’s weight in rats^10,20^ Habituation took place on the two days preceding the pretest. Actors and partners were placed in the assigned compartments for 10 minutes to freely explore all compartments. Reward was available *ad libitum*. During the pretest, only actors were present in the cage. The pretest session consisted of 6 forced choice trials and 16 free choice trials. The sequence of forced trials was always an alternation of right and left choices, starting with right.

At the beginning of each trial, the actor was placed in the starting compartment. Then, the doors were lifted, and the actor could access one of the reward compartments (during forced trials) or was offered a choice between the two compartments (during free choice trials). The actor received a food reward irrespective of choice (two chocolate chips, BioServ, 20 mg, #F05301). After a mouse consumed the reward, it was placed back in the starting compartment. The time mice could spend in each of the compartments was limited (**Fig. 1a**). In case the mouse did not move to the desired compartment before the time limit, the experimenter gently pushed it. The completion criterion for the pretest was 37 out of 44 food pellets consumed in two consecutive sessions (for the number of animals excluded based on this criterion, see **Figure S1** and **Table S1**). Additionally, an exclusion criterion was a >70% average preference for one of the compartments (the ‘70% criterion’). Number of animals that passed the predefined criteria is n=8 for females and n=10 for males.

During the main test phase, both the actor and partner were introduced to the cage, and testing sessions were performed daily over 4 days. Each actor’s entry to the “prosocial” compartment resulted in reward delivery for both mice. Conversely, upon entrance to the “asocial” compartment, only the actor was rewarded. The prosocial compartment was assigned as follows: in the case of mice with less than a 60% preference, the compartment was selected randomly. For the mice with an initial preference between 60 and 70%, the less preferred compartment was chosen as prosocial.

We considered the possibility that the prosocial behavior in female mice could be affected by estrous cycle phase. The estrous cycle in mice lasts for approximately four days. To minimize the possible effect of estrous cycle phase on the differences between males and females, the four-day average of the test phase results was used for pretest-posttest comparison and for comparison between sexes. The four day average was used for both sexes, to enable male-female comparison.

### Social conditioned place preference (sCPP)

sCPP was performed as previously described^37^,^46^. The procedure consisted of three phases: pretest, conditioning phase, and posttest.

During pretest each cage compartment contained one type of context (context A and context B) that differed in bedding type and gnawing block size and shape (**Table S13**). Both conditioning contexts were different from the home cage context, which consisted of aspen bedding (**Table S13**) and a distinct gnawing block. Mice were allowed to freely explore the cage for 30 minutes. Two animals were tested in parallel in adjacent cages. The exclusion criterion for pretest was initial preference to any of the context exceeding 70% (for the number of animals excluded based on this criterion, see **Table S14**). Number of animals that passed the predefined criteria is n=16 for females, n=12 for males tested in the 6-day protocol and n=8 for males tested in 2-day protocol.

After the pretest, animals were returned to their home cages for approximately 24 h. Then, mice were subjected to social conditioning (housing with cage mates) for 24 h in one of the contexts used in the pretest followed by 24 h of isolated conditioning (single housing) in the other context. To preserve an unbiased design, the social context was randomly assigned in such a way that approximately half of the cages received social conditioning on context A and half on context B. The conditioning phase lasted 6 days (3 days in each context, alternating every 24 h). After conditioning, the post-test was performed identically as pretest. Two sets of conditioning contexts were used (**Table S13**), and the results from both sets were pooled. When the number of animals conditioned on different bedding types (contexts) were not equal, the number of animals for each type of bedding was equalized by randomly removing the appropriate number of cases from the larger group using an R script as described in^37^.

### Affective state discrimination

The test was based on the protocol developed in 2020 by Scheggia and colleagues^26^. The behavior was assessed in a rectangular cage with opaque walls (see Fig. 3A; 53 cm × 32 cm × 15 cm). Demonstrators were placed on plastic platforms and confined under wire cups (diameter 9.5 cm × height 9 cm, Warmet, #B-0197). The procedure comprised two phases: habituation (3 days) and testing (1 day). The largest animal in the cage at the start of habituation was always assigned the “observer” role, the second largest was assigned the “relieved” or ”hungry” demonstrator role, and the smallest was assigned the “neutral” demonstrator. This was done to match the role assignment in the prosocial choice task and to ensure that the relieved/hungry and neutral demonstrators had a similar weight such that the only characteristic that distinguished them was the affective state. The relieved/hungry demonstrator and observer were always naïve, while the neutral demonstrators were tested twice with different observers in 9 cases, always a week apart.

On the first day of habituation, the observers were placed in the experimental cage for 12 minutes. In the experiment with relieved demonstrators the cage was empty for half of the animals, and it contained empty wire cups for the other half. No effect of cup presence during habituation was observed (**Fig. S3c-d**, **Table S15**). In the experiment with hungry demonstrators the cage always contained empty wire cups on the first day of habituation. On habituation days 2 and 3, observers were placed in the experimental cage for 6 minutes, and the wire cups were introduced for the next 6 minutes to habituate the observer to their placement during the test.

A glass jar was always placed on the top of the wired cups to prevent the observers from climbing the cups. Demonstrators were placed every day for 10 minutes in the experimental cage under the wired cups without an observer present. After the last habituation session, animals were placed in separate home cages for 24 hours, i.e., until the main test. The relieved/hungry demonstrators were deprived of food immediately after being put in a separate cage, while neutral demonstrators and observers had access to food *ad libitum*. One hour before the test, the relieved demonstrators were provided *ad libitum* access to food. Ten minutes before the test, observers were placed in the testing arena for habituation, and demonstrators were placed under wire cups on the table in the experimental room. Additionally, hungry demonstrators were presented with two chow pellets placed in unreachable distance to the wired cage to induce stress. After habituation, two demonstrators (neutral and relieved or hungry) were placed in the arena for 4 minutes (under wire cups). Observers who investigated both partners for less than 30 seconds were excluded from the analysis (for the number of animals excluded based on this criterion, see **Table S16**). The “relieved” and “hungry” conditions were tested in two consecutive experiments, on separate groups of animals. Number of animals that passed the predefined criteria for “relieved” experiment was n=12 for females and n=13 for males, and for “hungry” experiment: n = 8 for females and n=10 for males.

## Data analysis

Distance moved and presence in separate cage compartments in the sCPP test were automatically analyzed using EthoVision XT 15 software (Noldus, The Netherlands). Time spent sniffing relieved/hungry and neutral demonstrators and time spent in the respective zones were scored manually using Boris software^47^ by the experimenter, who was blinded to the demonstrators’ state. The significance level was set at p < 0.05. Comparisons of sample means were performed using analysis of variance (ANOVA) with Geisser-Greenhouse correction followed by Šídák’s multiple comparisons test or Student’s t-test for cases with only two samples.

## Acknowledgments

This work was supported by grant OPUS UMO-2016/21/B/NZ4/00198 from the National Science Centre Poland and the statutory funds of the Maj Institute of Pharmacology of the Polish Academy of Sciences. Special thanks to Katarzyna Kubik for help in analysis of the recordings from affective state discrimination experiment.

## Author contributions

KM, JRP and ZH designed the study. KM, ZH, MK, MC, ŁS, AB performed experiments. KM, KP and ZH analyzed the data. JRP and ZH supervised the study. JRP, ZH and KM wrote the paper with contributions from all of the authors.

## Data availability statement

All data are available at https://zenodo.org/record/6988393. Raw video recordings of the tests will be made available on request.

---

**Supplementa1 Figure S1.**
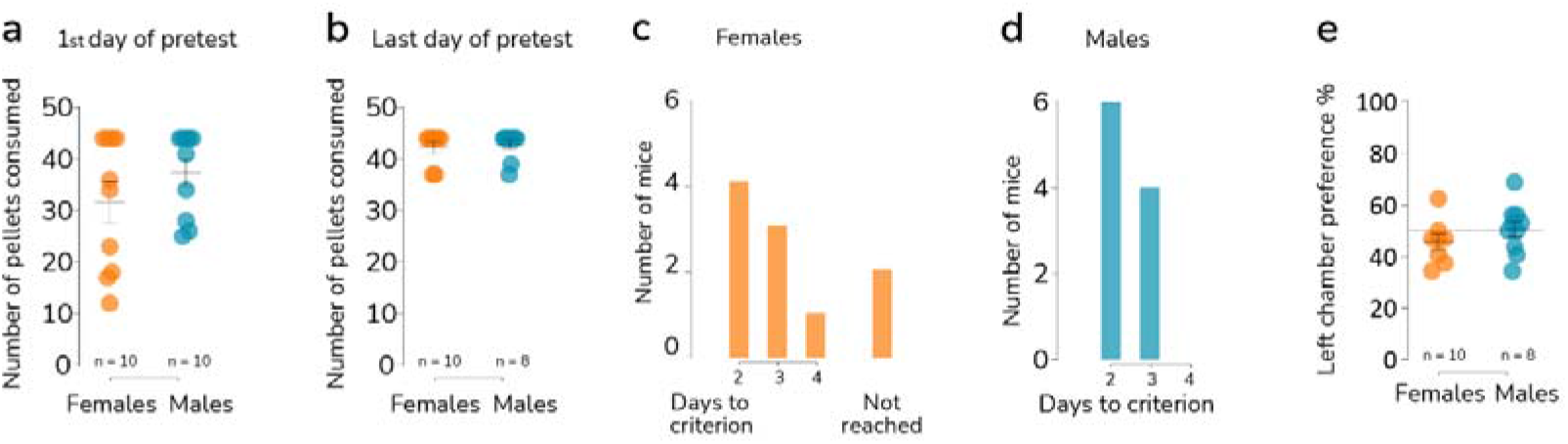
Rewards consumed during the pretest phase of the prosocial task. (**a-b**) The number of food pellets consumed by the female and male actors during the first and final days of the pretest, respectively. Each point represents an individual female or male mouse, with the mean and s.e.m. shown in black and the group sizes indicated below. (**c-d**) The number of female and male mice that reached the inclusion criterion of 37 out of 44 food pellets consumed on two consecutive sessions. As shown in (**c**), two female animals did not meet the required criterion. (**e**) No differences form chance level in average preference for the left chamber of the testing apparatus during pretest in female and male mice, respectively.

**Supplementa1 Figure S2.**
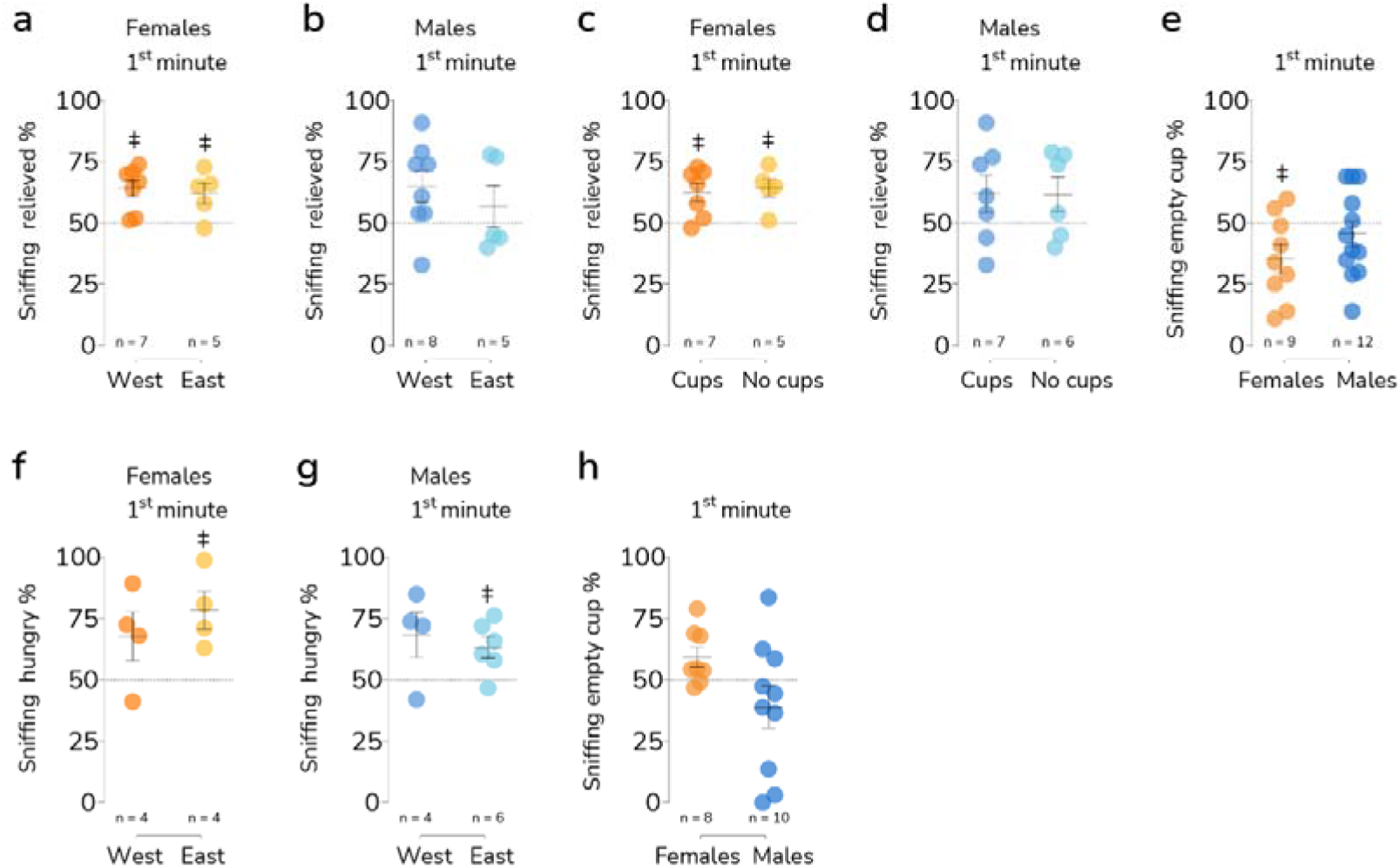
Context effects in the affective state determination task. (**a-b**) No effect of the position of the relieved demonstrator (West or East) on the proportion of the time the observer spent sniffing him in female and male mice, respectively. A significant preference (greater than 0) is indicated by a “‡” (one-sample t-test p<0.05) (**c-d**) No effect of the presence of the cups during habituation. (**e**) Final day of habituation. No effect of sex on time spent sniffing the empty cup in which the relieved demonstrator was placed during the test. Recordings of final day of habituation of 3 females and 1 male have been lost. (**f-g**) No effect of the position of the hungry demonstrator (West or East) on the proportion of the time the observer spent sniffing him in female and male mice, respectively. (**h**) Final day of habituation. No effect of sex on time spent sniffing the empty cup in which the hungry demonstrator was placed during the test.

**Table S1.**
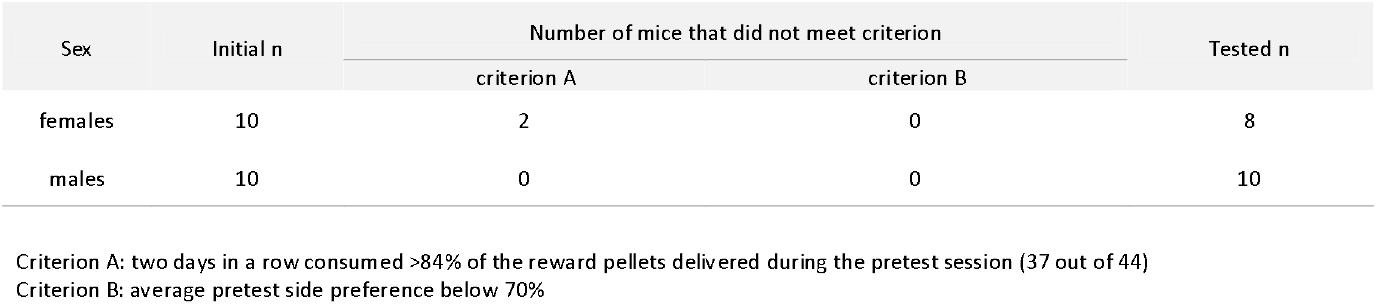
Prosocial choice test. Number of mice excluded based on predefined criteria.

**Table S2.** Prosocial choice test. Average pretest and test preference for prosocial compartment. **Table S2.**Average preference for the left chamber of the testing apparatus during pretest.

| Fig. | Sex | n | Preference during pretest % |  | Preference during test % |  | Mean difference | Paired Student's t-test |  |  |
| --- | --- | --- | --- | --- | --- | --- | --- | --- | --- | --- |
|  |  |  | M | SEM | M | SEM |  | t | df | p |
| 1e | females | 8 | 44,10 | 2,70 | 54,30 | 3,00 | 10,16 | 4,33 | 7,00 | 0,00 |
| 1f | males | 10 | 47,50 | 2,90 | 47,80 | 1,50 | 0,31 | 0,10 | 9,00 | 0,92 |

**Table S2. Continued.** Average preference for the left chamber of the testing apparatus during pretest.
| Fig. | Sex | n | M | SEM | One sample t test. Comparison with hypothetical mean (50%) |  |  | Mean difference | Unpaired Student's t-test |  |  |
| --- | --- | --- | --- | --- | --- | --- | --- | --- | --- | --- | --- |
|  |  |  |  |  | t | df | p |  | t | df | p |
| S1e | females | 8 | 45,71 | 3,07 | 1,40 | 7,00 | 0,20 | 4,93 | 1,13 | 16,00 | 0,27 |
|  | males | 10 | 50,64 | 3,015 | 0,21 | 9,00 | 0,84 |  |  |  |  |

**Table S3.** Prosocial choice test. Prosociality score. Females and males comparison.

| Fig. | Sex | Prosociality score %<sup>a</sup> |  |  |  |  |  |  |  |  |  |
| --- | --- | --- | --- | --- | --- | --- | --- | --- | --- | --- | --- |
|  |  | n | M | SEM | One sample t test. Comparison with hypothetical mean (0) |  |  | Mean difference | Unpaired Student's t-test |  |  |
|  |  |  |  |  | t | df | p |  | t | df | p |
| 1g | females | 8 | 10,20 | 2,30 | 4,35 | 7 | 0,00 | 9,88 | 2,47 | 16,00 | 0,03 |
|  | males | 10 | 0,30 | 3,00 | 0,10 | 9 | 0,92 |  |  |  |  |

<sup>a</sup> Mean % of prosocial choices of the last two days of pretest subtracted from mean % of prosocial choices of 4 days of test.

**Table S4.** Descriptive Statistics and Correlations.

| Sex | Behavioral test | Experiment variant | Variable | n | M | SEM | Subject's weight [g]<sup>a</sup> |  |  | Stimulus' weight [g]<sup>b</sup> |  |  | Weight difference %<sup>c</sup> |  |  |
| --- | --- | --- | --- | --- | --- | --- | --- | --- | --- | --- | --- | --- | --- | --- | --- |
|  |  |  |  |  |  |  | r<sup>2</sup> | r | p | r<sup>2</sup> | r | p | r<sup>2</sup> | r | p |
| females | Prosocial choice test<sup>1</sup> |  | Prosociality score % | 8 | 10,18 | 2,34 | 0,00 | 0,06 | 0,88 | 0,09 | 0,30 | 0,46 | 0,05 | -0,22 | 0,59 |
|  | Affective state discrimination test<sup>2</sup> | relieved | Sniffing relieved % | 12 | 63,83 | 2,72 | 0,40 | -0,62 | **0,02** | 0,50 | -0,70 | **0,00** | 0,28 | 0,53 | 0,07 |
|  |  | hungry | Sniffing hungry % | 8 | 73,10 | 6,18 | 0,19 | -0,44 | 0,28 | 0,00 | 0,04 | 0,92 | 0,48 | -0,69 | 0,06 |
|  | Social conditioned place preference test<sup>3</sup> |  | Preference score [s] | 16 | 285,80 | 87,07 | 0,09 | 0,31 | 0,24 | NA | NA | NA | NA | NA | NA |
| males | Prosocial choice test<sup>1</sup> |  | Prosociality score % | 10 | 0,30 | 3,03 | 0,13 | -0,36 | 0,38 | 0,09 | 0,30 | 0,30 | 0,54 | -0,73 | **0,01** |
|  | Affective state discrimination test<sup>2</sup> | relieved | Sniffing relieved % | 13 | 62,15 | 4,96 | 0,07 | 0,26 | 0,37 | 0,05 | -0,24 | 0,42 | 0,00 | 0,02 | 0,93 |
|  |  | hungry | Sniffing hungry % | 10 | 65,22 | 4,26 | 0,00 | -0,03 | 0,93 | 0,04 | -0,19 | 0,59 | 0,09 | 0,30 | 0,39 |
|  | Social conditioned place preference test<sup>3</sup> |  | Preference score [s] | 12 | 221,60 | 90,59 | 0,04 | -0,12 | 0,54 | NA | NA | NA | NA | NA | NA |

1a. Actor's weight on the first day of test; 1b. Partner's weight on the first day of test; 1c. Percentage of weight difference between actor and partner on the first day of test. 2a. Observer's weight on the first day of adaptation; 2b. Relieved/hungry demonstrator's weight on the first day of adaptation; 2c. Percentage of weight difference between observer and relieved/hungry demonstrator on the first day of adaptation. 3a. Subject's weight on the day of posttest.

**Table S5.** Social conditioned place preference test. Average pretest and test preference for social context.

| Fig. | Sex | n | Preference during pretest [s] |  | Preference during test [s] |  | Mean difference | Paired Student's t-test |  |  |
| --- | --- | --- | --- | --- | --- | --- | --- | --- | --- | --- |
|  |  |  | M | SEM | M | SEM |  | t | df | p |
| 2a | females | 16 | 893,70 | 34,85 | 1040,00 | 43,56 | 146,30 | 2,83 | 15,00 | **0,01** |
| 2b | males | 12 | 855,80 | 40,59 | 1010,00 | 45,28 | 154,20 | 4,20 | 11,00 | **0,00** |
| 2d | males | 8 | 918,90 | 26,18 | 838,20 | 58,76 | 80,69 | 1,18 | 7,00 | 0,28 |

**Table S6.** Social conditioned place preference test. Preference score. Females and males comparison.

| Fig. | Sex | n | M | SEM | Social preference score [s]<sup>a</sup> |  |  |  |  |  |  |
| --- | --- | --- | --- | --- | --- | --- | --- | --- | --- | --- | --- |
|  |  |  |  |  | One sample t test. Comparison with hypothetical mean (0) |  |  | Mean difference | Unpaired Student's t-test |  |  |
|  |  |  |  |  | t | df | p |  | t | df | p |
| 2c | females | 16 | 285,80 | 87,07 | 3,28 | 15,00 | **0,01** | 64,20 | 0,50 | 26,00 | 0,62 |
|  | males | 12 | 221,60 | 90,59 | 2,45 | 11,00 | **0,03** |  |  |  |  |
| 2d | males | 8 | -119,40 | 117,70 | 1,02 | 7,00 | 0,34 |  |  |  |  |

<sup>a</sup> Time spent in social context pretest [s] subtracted from time spent in social context posttest [s]

**Table S7.** Affective state discrimination test. Anova with repeated measures table for sniffing demonstrators [s].

| Demonstrators state | Fig. | Sex | Source of variation |  |  |  |  |  |  |  |  |  |  |  |  |  |  |
| --- | --- | --- | --- | --- | --- | --- | --- | --- | --- | --- | --- | --- | --- | --- | --- | --- | --- |
|  |  |  | Time |  |  |  |  | Demonstrator's state |  |  |  |  | Interaction |  |  |  |  |
|  |  |  | SS | DF | MS | F | p | SS | DF | MS | F | p | SS | DF | MS | F | p |
| relieved | 3b | females | 315,50 | 3,00 | 105,20 | 3,68 | 0,02 | 106,90 | 1,00 | 106,90 | 2,33 | 0,14 | 208,30 | 3,00 | 69,44 | 2,43 | 0,07 |
|  | 3c | males | 129,50 | 3,00 | 43,16 | 1,25 | 0,30 | 584,30 | 1,00 | 584,30 | 6,23 | 0,02 | 185,90 | 3,00 | 61,98 | 1,80 | 0,15 |
| hungry | 3e | females | 49,48 | 3,00 | 16,49 | 0,60 | 0,57 | 370,30 | 1,00 | 370,30 | 5,75 | 0,03 | 425,00 | 3,00 | 141,70 | 5,16 | 0,00 |
|  | 3f | males | 323,10 | 3,00 | 107,70 | 5,53 | 0,00 | 45,52 | 1,00 | 45,52 | 0,80 | 0,38 | 314,80 | 3,00 | 104,90 | 5,39 | 0,00 |

**Table S7. Continued.**
| Demonstrators state | Fig. | Sex | Comparison of time sniffing relieved vs neutral demonstrators [s] |
| --- | --- | --- | --- |

|  |  |  | Time [minute] | M<sub>relieved</sub> | M<sub>neutral</sub> | Mean difference | Šidák's multiple comparisons test |  |  |  |
| --- | --- | --- | --- | --- | --- | --- | --- | --- | --- | --- |
|  |  |  |  |  |  |  | SE | t | DF | p |
| relieved | 3b | females | 1<sup>st</sup> | 17,53 | 10,46 | 7,07 | 2,25 | 3,14 | 21,74 | **0,02** |
|  |  |  | 2<sup>nd</sup> | 13,83 | 12,32 | 1,52 | 2,41 | 0,63 | 21,19 | 0,95 |
|  |  |  | 3<sup>rd</sup> | 12,53 | 12,22 | 0,31 | 2,57 | 0,12 | 21,96 | 1,00 |
|  |  |  | 4<sup>th</sup> | 8,95 | 9,40 | -0,45 | 2,12 | 0,21 | 20,55 | 1,00 |
|  | 3c | males | 1<sup>st</sup> | 20,25 | 12,17 | 8,09 | 2,61 | 3,09 | 23,59 | **0,02** |
|  |  |  | 2<sup>nd</sup> | 18,09 | 12,51 | 5,59 | 2,77 | 2,01 | 19,25 | 0,21 |
|  |  |  | 3<sup>rd</sup> | 14,44 | 13,78 | 0,65 | 2,78 | 0,24 | 18,91 | 1,00 |
|  |  |  | 4<sup>th</sup> | 15,61 | 10,97 | 4,64 | 2,84 | 1,64 | 19,79 | 0,39 |
| hungry | 3e | females | 1<sup>st</sup> | 17,44 | 5,92 | 11,53 | 2,81 | 4,10 | 10,62 | **0,01** |
|  |  |  | 2<sup>nd</sup> | 13,10 | 7,96 | 5,14 | 2,97 | 1,73 | 11,98 | 0,37 |
|  |  |  | 3<sup>rd</sup> | 7,72 | 10,69 | -2,97 | 3,46 | 0,86 | 12,37 | 0,88 |
|  |  |  | 4<sup>th</sup> | 13,09 | 7,55 | 5,54 | 2,84 | 1,95 | 13,92 | 0,26 |
|  | 3f | males | 1<sup>st</sup> | 16,34 | 8,93 | 7,41 | 2,40 | 3,08 | 17,21 | **0,03** |
|  |  |  | 2<sup>nd</sup> | 11,88 | 9,33 | 2,55 | 2,17 | 1,18 | 17,98 | 0,69 |
|  |  |  | 3<sup>rd</sup> | 7,79 | 10,98 | -3,19 | 2,78 | 1,15 | 16,55 | 0,71 |
|  |  |  | 4<sup>th</sup> | 6,72 | 7,45 | -0,73 | 2,20 | 0,33 | 17,99 | 1,00 |

**Table S8.**
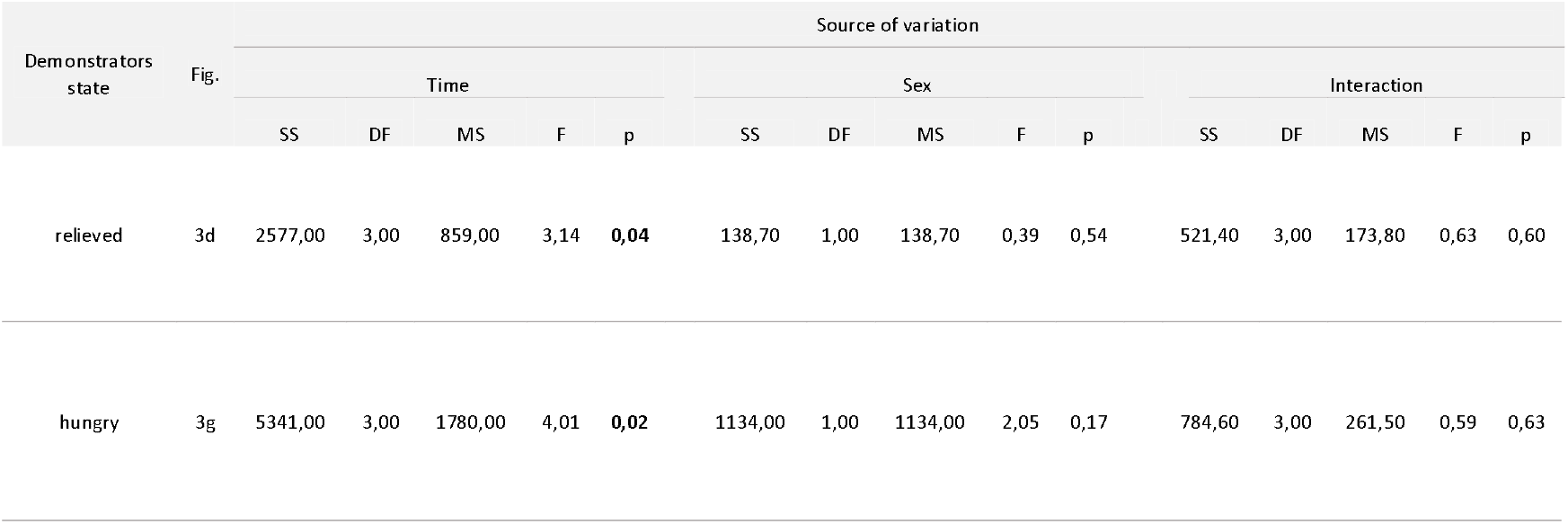
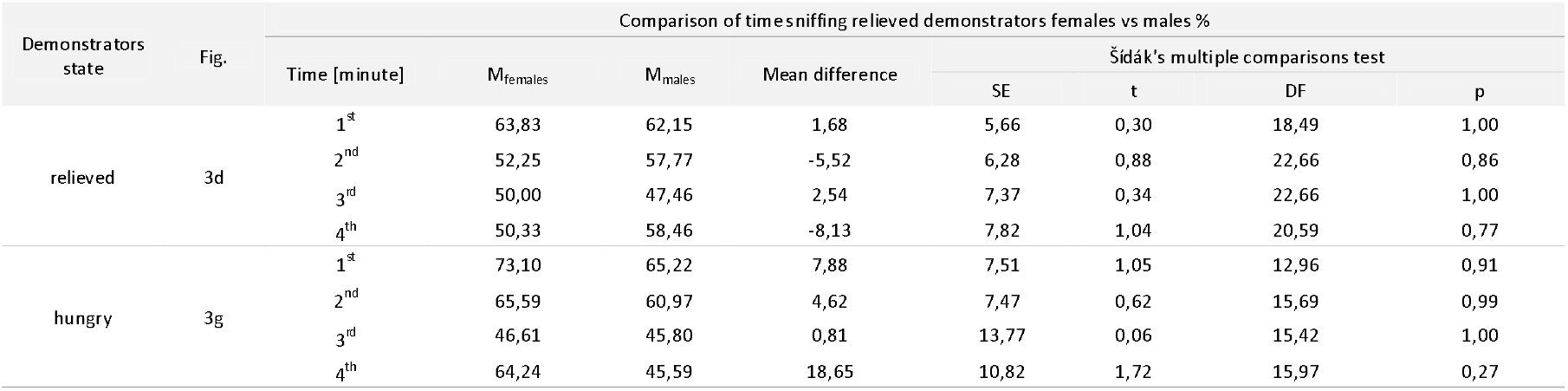
Affective state discrimination test. Anova with repeated measures table for sniffing demonstrators %

**Table S9.** Affective state discrimination test. One sample t-test for sniffing relieved demonstrators %

| Demonstrators state | Fig. | Sex | Sniffing relieved/hungry demonstrators % |  |  |  |  |  |  |
| --- | --- | --- | --- | --- | --- | --- | --- | --- | --- |
|  |  |  | n | M | SEM | Time [minute] | One sample t test. Comparison with hypothetical mean (50%) |  |  |
|  |  |  |  |  |  |  | t | df | p |
| relieved | 3d | females | 12 | 63,83 | 2,72 | 1<sup>st</sup> | 5,08 | 11,00 | **0,00** |
|  |  |  |  | 52,25 | 4,61 | 2<sup>nd</sup> | 0,49 | 11,00 | 0,64 |
|  |  |  |  | 50,00 | 4,76 | 3<sup>rd</sup> | 0,00 | 11,00 | 1,00 |
|  |  |  |  | 50,33 | 6,30 | 4<sup>th</sup> | 0,05 | 11,00 | 0,96 |
|  |  | males | 13 | 62,15 | 4,96 | 1<sup>st</sup> | 2,45 | 13,00 | **0,03** |
|  |  |  |  | 57,77 | 4,265 | 2<sup>nd</sup> | 1,82 | 13,00 | 0,09 |
|  |  |  |  | 47,46 | 5,629 | 3<sup>rd</sup> | 0,45 | 13,00 | 0,66 |
|  |  |  |  | 58,46 | 4,635 | 4<sup>th</sup> | 1,83 | 13,00 | 0,09 |
| hungry | 3g | females | 8 | 73,10 | 6,18 | 1<sup>st</sup> | 3,74 | 7,00 | **0,01** |
|  |  |  |  | 65,59 | 5,32 | 2<sup>nd</sup> | 2,93 | 7,00 | **0,02** |
|  |  |  |  | 46,61 | 10,05 | 3<sup>rd</sup> | 0,34 | 7,00 | 0,75 |
|  |  |  |  | 64,24 | 7,33 | 4<sup>th</sup> | 1,94 | 7,00 | 0,09 |
|  |  | males | 10 | 65,22 | 4,261 | 1<sup>st</sup> | 3,57 | 9,00 | **0,01** |
|  |  |  |  | 60,97 | 5,238 | 2<sup>nd</sup> | 2,09 | 9,00 | 0,07 |
|  |  |  |  | 45,80 | 9,404 | 3<sup>rd</sup> | 0,45 | 9,00 | 0,67 |
|  |  |  |  | 45,59 | 7,961 | 4<sup>th</sup> | 0,55 | 9,00 | 0,59 |

**Table S10.** Affective state discrimination test. One sample t-test for sniffing relieved demonstrators %. Demonstrators position (east vs west) comparison.

| Demonstrators state | Fig. | Sex | Demonstrator's position | Sniffing relieved/hungry demonstrators % |  |  |  |  |  |  |  |  |  |
| --- | --- | --- | --- | --- | --- | --- | --- | --- | --- | --- | --- | --- | --- |
|  |  |  |  | n | M | SEM | One sample t test. Comparison with hypothetical mean (50%) |  |  | Mean difference | Unpaired Student's t-test |  |  |
|  |  |  |  |  |  |  | t | df | p |  | t | df | p |
| relieved | S2a | females | west | 7 | 64,14 | 3,47 | 4,07 | 6,00 | **0,01** | 2,14 | 0,39 | 10,00 | 0,70 |
|  |  |  | east | 5 | 62,00 | 4,23 | 2,84 | 4,00 | **0,05** |  |  |  |  |
|  | S2b | males | west | 8 | 65,00 | 6,43 | 2,33 | 7,00 | 0,05 | 8,20 | 0,78 | 11,00 | 0,45 |
|  |  |  | east | 5 | 56,80 | 8,49 | 0,80 | 4,00 | 0,47 |  |  |  |  |
| hungry | S2f | females | west | 4 | 67,76 | 9,97 | 1,78 | 3,00 | 0,17 | 10,67 | 0,85 | 6,00 | 0,43 |
|  |  |  | east | 4 | 78,43 | 7,74 | 3,67 | 3,00 | **0,03** |  |  |  |  |
|  | S2g | males | west | 4 | 68,24 | 9,20 | 1,98 | 3,00 | 0,14 | 5,03 | 0,56 | 8,00 | 0,59 |
|  |  |  | east | 6 | 63,21 | 4,32 | 3,06 | 5,00 | **0,03** |  |  |  |  |

**Table S11.** Affective state discrimination test. One sample t-test for sniffing relieved/hungry to be demonstrators %.

| Demonstrators state | Fig. | Sex | Sniffing relieved/hungry to be demonstrators % |  |  |  |  |  |  |  |  |  |
| --- | --- | --- | --- | --- | --- | --- | --- | --- | --- | --- | --- | --- |
|  |  |  | n | M | SEM | One sample t test. Comparison with hypothetical mean (50%) |  |  | Mean difference | Unpaired Student's t-test |  |  |
|  |  |  |  |  |  | t | df | p |  | t | df | p |
| relieved | S2e | females | 9 | 35,44 | 5,83 | 2,50 | 8,00 | **0,04** | 10,06 | 1,28 | 19,00 | 0,22 |
|  |  | males | 12 | 45,50 | 5,19 | 0,87 | 11,00 | 0,40 |  |  |  |  |
| hungry | S2h | females | 8 | 59,32 | 3,98 | 2,34 | 7,00 | 0,05 | 20,46 | 2,01 | 16,00 | 0,06 |
|  |  | males | 10 | 38,86 | 8,50 | 1,31 | 9,00 | 0,22 |  |  |  |  |

**Table S12.** Groups of animals used in Figures 1-3. Only mice that met predefined criteria.

| Sex | Behavioral test | Experiment variant | n | Subject's age at the start of the procedure (weeks)<sup>a</sup> |  |  | Subject's age at test (weeks)<sup>b</sup> |  |  | Subject's weight at the start of the procedure [g]<sup>a</sup> |  |  |
| --- | --- | --- | --- | --- | --- | --- | --- | --- | --- | --- | --- | --- |
|  |  |  |  | M | Range | SEM | M | Range | SEM | M | Range | SEM |
| females | Prosocial choice test<sup>1</sup> |  | 8 | 10,30 | 9,90-10,40 | 0,06 | 11,70 | 11,30-11,90 | 0,07 | 19,85 | 18,60-21,00 | 0,28 |
|  | Affective state discrimination test<sup>2</sup> | relieved | 12 | 12,62 | 10,90-14,60 | 0,31 | 13,09 | 11,30-15,90 | 0,36 | 20,38 | 18,50-22,20 | 0,37 |
|  |  | hungry | 8 | 11,49 | 9,90-13,40 | 0,52 | 11,94 | 10,30-13,90 | 0,53 | 21,28 | 19,40-23,20 | 0,41 |
|  | Social conditioned place preference test<sup>3</sup> |  | 16 | 10,73 | 10,60-10,90 | 0,03 | 11,56 | 11,40-11,90 | 0,05 | 18,69 | 17,40-19,60 | 0,28 |
| males | Prosocial choice test<sup>1</sup> |  | 10 | 10,30 | 9,90-11,00 | 0,14 | 11,70 | 11,00-12,40 | 0,15 | 26,65 | 24,10-28,60 | 0,45 |
|  | Affective state discrimination test<sup>2</sup> | relieved | 13 | 11,40 | 9,90-12,60 | 0,24 | 11,95 | 10,30-14,00 | 0,30 | 26,05 | 23,30-29,10 | 0,49 |
|  |  | hungry | 10 | 11,52 | 10,00-14,10 | 0,43 | 12,07 | 10,70-14,60 | 0,43 | 26,69 | 24,80-31,00 | 0,58 |
|  | Social conditioned place preference test<sup>3</sup> |  | 12 | 10,60 | 10,00-11,00 | 0,14 | 11,60 | 11,00-12,00 | 0,14 | 23,90 | 21,30-26,00 | 0,40 |

**Table S12.** Continued.
| Sex | Behavioral test | Experiment variant | n | Subject's weight at test [g]<sup>b</sup> |  |  | Stimulus' weight at the start of the procedure [g]<sup>c</sup> |  |  | Stimulus' weight at test [g]<sup>d</sup> |  |  |
| --- | --- | --- | --- | --- | --- | --- | --- | --- | --- | --- | --- | --- |
|  |  |  |  | M | Range | SEM | M | Range | SEM | M | Range | SEM |
| females | Prosocial choice test<sup>1</sup> |  | 8 | 17,01 | 15,80-18,40 | 0,34 | 18,25 | 16,70-19,90 | 0,34 | 15,50 | 13,8-16,90 | 0,34 |
|  | Affective state discrimination test<sup>2</sup> | relieved | 12 | 19,83 | 17,20-21,50 | 0,62 | 19,68 | 17,40-21,30 | 0,42 | 16,53 | 15,40-18,70 | 0,49 |
|  |  | hungry | 8 | 21,70 | 21,20-22,50 | 0,22 | 20,65 | 19,40-21,60 | 0,25 | 17,68 | 17,20-18,30 | 0,20 |
|  | Social conditioned place preference test<sup>3</sup> |  | 16 | 19,82 | 17,40-21,50 | 0,31 | NA | NA | NA | NA | NA | NA |
| males | Prosocial choice test<sup>1</sup> |  | 10 | 22,81 | 20,60-25,40 | 0,47 | 25,24 | 20,60-25,40 | 0,60 | 21,40 | 22,90-27,40 | 0,45 |
|  | Affective state discrimination test<sup>2</sup> | relieved | 13 | 25,42 | 23,20-28,10 | 0,51 | 24,65 | 21,00-28,20 | 0,57 | 20,95 | 17,30-26,80 | 0,73 |
|  |  | hungry | 10 | 27,23 | 24,80-30,20 | 0,74 | 25,21 | 23,40-28,60 | 0,49 | 21,56 | 19,90-23,10 | 0,65 |
|  | Social conditioned place preference test<sup>3</sup> |  | 12 | 23,74 | 21,60-25,50 | 0,34 | NA | NA | NA | NA | NA | NA |

**1a.** Actor's age/weight at start of the food deprivation; **1b.** Actor's age/weight on the first day of test; **1c.** Partner's weight on the start of the food deprivation; **1d.** Partner's weight on the first day of test. **2a.** Observer's age/weight on the first day of adaptation; **2b.** Observer's age/weight on test day; **2c.** Relieved/hungry demonstrator's weight on the first day of adaptation. **3a.** Subject's age/weight on the day of pretest; **3b.** Subject's age/weight on the day of posttest.

**Table S13.**
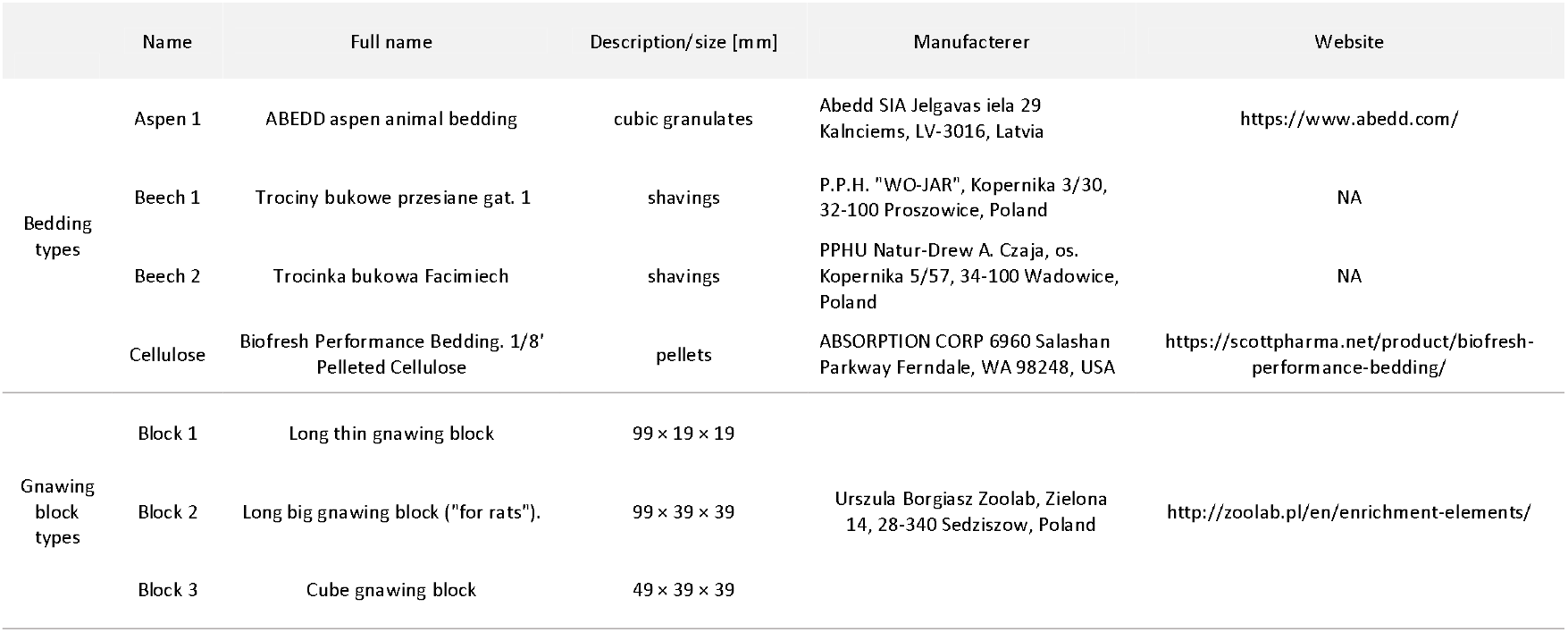
Social conditioned place preference test. Conditioning cues.

**Table S14.** Social conditioned place preference test. Number of mice excluded based on predefined criteria.

| sCPP protocol | Sex | Initial n | Number of mice that did not meet criterion A | Tested n |
| --- | --- | --- | --- | --- |
| 6 days | females | 17 | 1 | 16 |
| 6 days | males | 12 | 0 | 12 |
| 2 days | males | 8 | 0 | 8 |

Criterion A: Initial preference to any of the context not exceeding 70% in pretest.

**Table S15.** Affective state discrimination test. One sample t-test for sniffing relieved demonstrators %. Demonstrators position (cups vs no cups) comparison.

| Demonstrators state | Fig. | Sex | Demonstrator's position | Sniffing relieved/hungry demonstrators % |  |  |  |  |  |  |  |  |  |
| --- | --- | --- | --- | --- | --- | --- | --- | --- | --- | --- | --- | --- | --- |
|  |  |  |  | n | M | SEM | One sample t test.<br>Comparison with<br>hypothetical mean (50%) |  |  | Mean difference | Unpaired Student's t-test |  |  |
|  |  |  |  |  |  |  | t | df | p |  | t | df | p |
| relieved | S2c | females | cups | 7 | 62,57 | 3,75 | 3,35 | 6,00 | 0,02 | 1,63 | 0,30 | 10,00 | 0,77 |
|  |  |  | no cups | 5 | 64,20 | 3,73 | 3,80 | 4,00 | 0,02 |  |  |  |  |
|  | S2d | males | cups | 7 | 62,00 | 7,62 | 1,57 | 6,00 | 0,17 | 0,33 | 0,03 | 11,00 | 0,98 |
|  |  |  | no cups | 6 | 61,67 | 7,13 | 1,64 | 5,00 | 0,16 |  |  |  |  |

In the experiments with hungry demonstrators during the first day of habituation there were always cups in test cage.

**Table S16.** Affective state discrimination test. Number of mice excluded based on predefined criteria.

| Demonstrators state | Sex | Initial n | Number of mice that did not meet criterion A | Tested n |
| --- | --- | --- | --- | --- |
| relieved | females | 14 | 2 | 12 |
|  | males | 15 | 2 | 13 |
| hungry | females | 9 | 1 | 8 |
|  | males | 11 | 1 | 10 |

